# Identification of potential new vaccine candidates in *Salmonella typhi* using reverse vaccinology and subtractive genomics-based approach

**DOI:** 10.1101/521518

**Authors:** Sumit Mukherjee, Kaustav Gangopadhyay, Sunanda Biswas Mukherjee

## Abstract

*Salmonella enterica serovar typhi* is the causative agent of typhoid fever in human. The available vaccine lacks the effectiveness and further overuse of the antibiotics throughout the past decades lead to the emergence of antibiotic-resistant strains. To reduce the spread of antibiotic resistance, it is essential to develop new vaccines. In this study using an extensive bioinformatic analysis based on reverse vaccinology and subtractive genomics-based approach, we searched the genome of *S typhi*. We found that two outermost membrane proteins LptD and LptE, which are responsible for the final transport of the LPS across the membrane having most of the essential signature for being vaccine target. Both of these proteins are found highly conserved and possess shared surface epitopes among several species of *Salmonella* pathogens. This indicates that these two proteins could be the potential target for vaccine development, which could be effective for a broad range of *Salmonella* pathogens.

## 1. Introduction

*Salmonella enterica serovar typhi* (*S typhi*) is a gram-negative human-restricted pathogenic bacterium that is the causative agent of an acute febrile illness known as typhoid fever. The World Health Organization (WHO) recognizes infection by *S typhi* as a persistent threat in developing and underdeveloped countries. The CDC reported that, globally, 22 million people affected by *S typhi* annually, resulted in approximately 600,000 deaths [1]. Mechanism of the pathogenesis of *S typhi* is still not completely understood. Consequent use of broad-spectrum antibiotics revealed the emergence of multi-drug resistant strains which is a major public health concern [2]. Vaccination is the most effective way, which can help to reduce the spread of antibiotic resistance. The resistance of the vaccine is very rare because pathogen populations create less variation for vaccine resistance than they do for drug resistance. The main underlying principles behind it that vaccines tend to work prophylactically by inducing immune responses against multiple pathogen proteins (antigens), and various epitopes on each antigen while drugs tend to work therapeutically by interfering with a specific step in a particular metabolic pathway [3]. At present, mainly two vaccines are available against typhoid fever which includes inactivated whole cell vaccines and live attenuated vaccines [4][5]. All of these vaccines have less than 80% efficacy [6][7]. Therefore, the development of new vaccines is necessary. Identification of potential vaccine target is the first step towards the development of a new vaccine. The advent of high–throughput sequencing technology provided a vast amount of genome sequences data of pathogens, which can be mined to explore the new potential vaccine targets. Reverse vaccinology is an *in-silico* approach that intended to detect potential vaccine protein targets from the genome of the pathogenic organisms [8][9]. Reverse vaccinology was first applied to pathogen *Neisseria meningitides* serogroup B (MenB) to identify vaccine candidates [10]. The vaccine deriving from the mining of the genome of *Neisseria meningitides* serogroup B was entered into the phase I clinical testing in 2002 after successful preclinical studies [11]. In November 2012, the first vaccine named Bexsero against meningococcal serogroup B was granted for marketing by European Medicinal Agency and a few months later in January 2013, it was also approved by the European Commission [12]. In the meantime, reverse vaccinology based approach has been applied to mining genome of pathogens such as *Bacillus anthracis* [13], *Streptococcus pneumonia [14], Mycobacterium tuberculosis [15], Cryptosporidium hominis [16], Helicobacter pyroli* [17] and *Treponema pallidum* [18].

In this present study, we have developed reverse vaccinology and subtractive genomics based computational strategy and screened the genome of *S typhi* and detected two highly potential vaccine candidates those contain most of the signatures of the ideal vaccine. All of the three proteins detected using this approach are essential for bacterial survival, extracellular membrane proteins and involved in the secretory pathway, containing the virulence determining factors and do not have any human homologous. Antigenic epitopes are present in those three proteins that have an affinity to bind B-cells and T-cells. Protein-protein interaction analysis reveals that those proteins actively interact with each other and all of them are involved in the LPS transport. Therefore, our findings put forward a set of new potential vaccine candidates of *S typhi*. This work can act as a benchmark and provide valuable input for further experimental and clinical testing.

## 2. Materials and Methods

### 2.1. Dataset collection

We retrieved the protein coding sequences of four strains (NC_003198.1, NC_016832.1, NC_021176.1, and NZ_CP012091.1) of *Salmonella enterica serovar typhi* from RefSeq database. We have performed the pan-genome analysis and those genes which are present in all of the four strains of *S typhi* were considered for our studies.

### 2.2. Selection of essential proteins

Essential proteins of pathogens are crucial for major functions for their survival within the host. Database of Essential Genes (DEG) [19] consists of the experimentally validated data for the essential genes within a genome. To identify the essential genes we performed BLASTP [20] searches of the protein sequences of all the strains of *S typhi* against DEG databases. The parameters for considering the essential genes were set as follow: E-value□≤□10-5, similarity □≥□ 50 %, and the sequence coverage□≤□30 %. Protein sequences of *S typhi* those have significant sequence similarity with the DEG protein sequences were retrieved.

### 2.3. Search for virulence factors in *S typhi*

Virulence factors are molecules synthesized by pathogens that contribute to the pathogenicity to infect the host. We predicted the pathogenic ability of the *S typhi* essential proteins using MP3 server (http://metagenomics.iiserb.ac.in/mp3/index.php) [21] (Gupta et al., 2014). This web-server predicts pathogenic and virulent proteins using an integrated SVM-HMM approach with high accuracy.

### 2.4. Secretory pathway analysis

Bacterial pathogens exploit diverse methods to invade mammalian hosts. One of the important strategies for many pathogens is the secretion of proteins across phospholipid membranes. Secreted proteins can take part in crucial roles in promoting bacterial virulence, by facilitating attachment to host cells and disrupting their functions [22]. SignalP web-server [23] was used to predict the signal peptide triggered protein secretion and SecretomeP 2.0 web-server [24] was used for predictions of non-classical i.e. not signal peptide triggered protein secretion.

### 2.5. Prediction of protein subcellular localization

Subcellular localization of *S typhi* proteins was predicted using two different web servers: CELLO v.2.5 (http://cello.life.nctu.edu.tw/) [25] and SubCellProt (http://www.databases.niper.ac.in/SubCellProt) [26]. CELLO predicts protein subcellular localization using two-level support vector machine (SVM). While, SubCellProt is based on two machine learning approaches, k Nearest Neighbor (k-NN) and Probabilistic Neural Network (PNN). When at least two of these three approaches (k-NN, PNN, and SVM) predict the same localization of a protein we consider that as its subcellular localization.

### 2.6. Screening of non-homologous proteins

To identify non-homologous proteins of *S typhi*, we used a systematic approach based on homology search. BLASTP [20] followed by TBLASTN filtering approach (E<10^−5^ and use of low-complexity filters) was used against all the human proteins to detect the *S typhi* proteins which have no homologs with humans.

### 2.7. Prediction of potential antigenic epitopes

The potentiality of the peptides to activate the human immune system are identified in order to generate memory cells against *S typhi*. To understand the immunomodulatory effect of these peptides, we have analyzed different types of epitopes. LBtope [27] was used to predict the linear B-cell epitopes. We used the LBtope model [27] with default threshold of 60% probability cut-off. Conformational B-cell epitopes were predicted using CBTOPE [28]. The major role of the MHC is to bind to antigenic regions (or peptides) and present these peptides on the cell surface, where appropriate T-cells recognize these peptides. Thus, it is important to predict MHC binders, as these binders have the ability to activate the T-cells of the immune system. nHLAPred [29] was used to predict MHC class-I binding peptide. In addition to the prediction of MHC binders which are potential T-cell epitopes, we also predicted CTL epitopes using a direct method, CTLPred [30]. CTLPred [30] is a direct method that predicts T-cell epitopes (CTL) from a primary sequence of the antigen instead of using the intermediate step where MHC Class I binders are predicted.

### 2.8. Molecular weight calculation

Protein molecular weight calculated using ExPasypI/Mw tool [31].

### 2.9. Protein interaction network analysis

A protein-protein interaction network of the selected proteins was constructed using the STRING database [32]. In the present study, all interactors with low confidence scores i.e., less than 0.400 were screened in order to avoid false positives and false negatives. Only medium and high confidence interactions with not more than 50 interactors were included in the interaction network. The protein networks constructed for selected proteins were then analyzed using Cytoscape 2.8.1 [33]. In order to evaluate the importance of the query node, each node in the protein network was deleted one by one and the change in clustering coefficient was analyzed [34][35]. Pathogen-specific essential proteins important in the metabolic network were regarded as potential targets.

## 3. Results

Reverse vaccinology is a promising approach for genome-wide identification of potential vaccine candidates. Systematically genome-wide screening is crucial for the identification of vaccine candidates from pathogens. We developed a systematic extensive computational screening approach based on reverse vaccinology and subtractive genomics for the identification of potential vaccine candidates from *S typhi*. Most of the important characteristic features that required for being potential vaccine candidates were considered for our analysis. The detailed computational pipelines are described in the methods section. Figure 1 represents the flow diagram of our detailed screening approaches and computational pipelines. For the initial screening step, we have selected only the essential genes of *S typhi*. Essential genes are involved in basic metabolic pathways and are crucial for the survival of any organisms. Therefore, in microbial pathogens, those genes constitute attractive targets for antimicrobial drug and vaccine design [36][37][38][39]. After the first step of screening, we have found total 358 genes, which are essential for *S typhi*. In the second step from the essential genes datasets, we have screened those genes, which comprise potential virulence factors. Virulence factors facilitate the pathogens to invade the host and evade host defenses. Genes having virulence factor induces pathogenicity within hosts and therefore, those genes can be a potential target for vaccines [40]. We have found only 20 proteins from essential genes datasets having significant pathogenic abilities. In the next step, we have searched the secreted proteins. Secreted proteins have vital roles in promoting bacterial virulence, from enhancing attachment to eukaryotic cells to scavenging resources in an environmental niche, to directly intoxicating target cells and disrupting their functions [22]. Therefore, protein secretion systems have an impact on bacterial pathogenesis and secretory proteins are ideal for vaccine targets. From the set of 20 protein sequences, we have predicted that 11 proteins have a role in the secretory pathway. In the next screening steps, we have select the extracellular membrane proteins. Membrane proteins play a key role in various bacterial physiology and pathogenesis and are the most important targets for vaccine development. We have analyzed the previously screened 11 proteins and predicted that 4 proteins are membrane proteins. Pathogen proteins those are not present in the humans are significant for vaccine targets because targeting inhibitors against these non-homologous sequences would have minimal side effects in humans [17][41]. Therefore, we have screened the protein sequences, which are non-homologous to human. From previously screened 5 protein datasets we have found 3 proteins which do not have sequence similarity with humans. Proteins having molecular weight <110 kDa are considered to be more effective targets because proteins having lower molecular weight can easily be purified and efficiently subjected to vaccine development [42]. We have found all of these three proteins have a molecular weight less than <100 kDa. Interacting partners of a protein at the cellular level can reveal many important signaling pathway-stimulating factors and some crucial cross-talks between potential therapeutic targets. Our protein interaction analysis revealed all these three proteins interact with each other and involved in bacterial LPS transport. The topological analysis of the constructed network indicate that all these three proteins interacts with other essential proteins involved in many important pathways for pathogen survival. In order to evaluate the importance of the each of these selected proteins, each protein in the network was deleted one by one and the change in clustering coefficient was analyzed. We found that all of these three proteins are crucial and deleting one node change the clustering coefficient. Next, we analyzed these three proteins to find the potential epitopes. We predicted that LptD and LptE both of these proteins contain multiple epitopes, and each of these protein contains one such epitopes each which can recognize B-cell, T-cell and can bind the MHC class-I. The epitopes are highlighted in the structure of LptDE complex and it has been observed that the epitopes in LptD and LptE are present in close proximity (Figure 2). These epitopes have high VaxiJen score (provided in Table 1) indicate that LptD and LptE could be the potential vaccine target.

**Figure 1:**
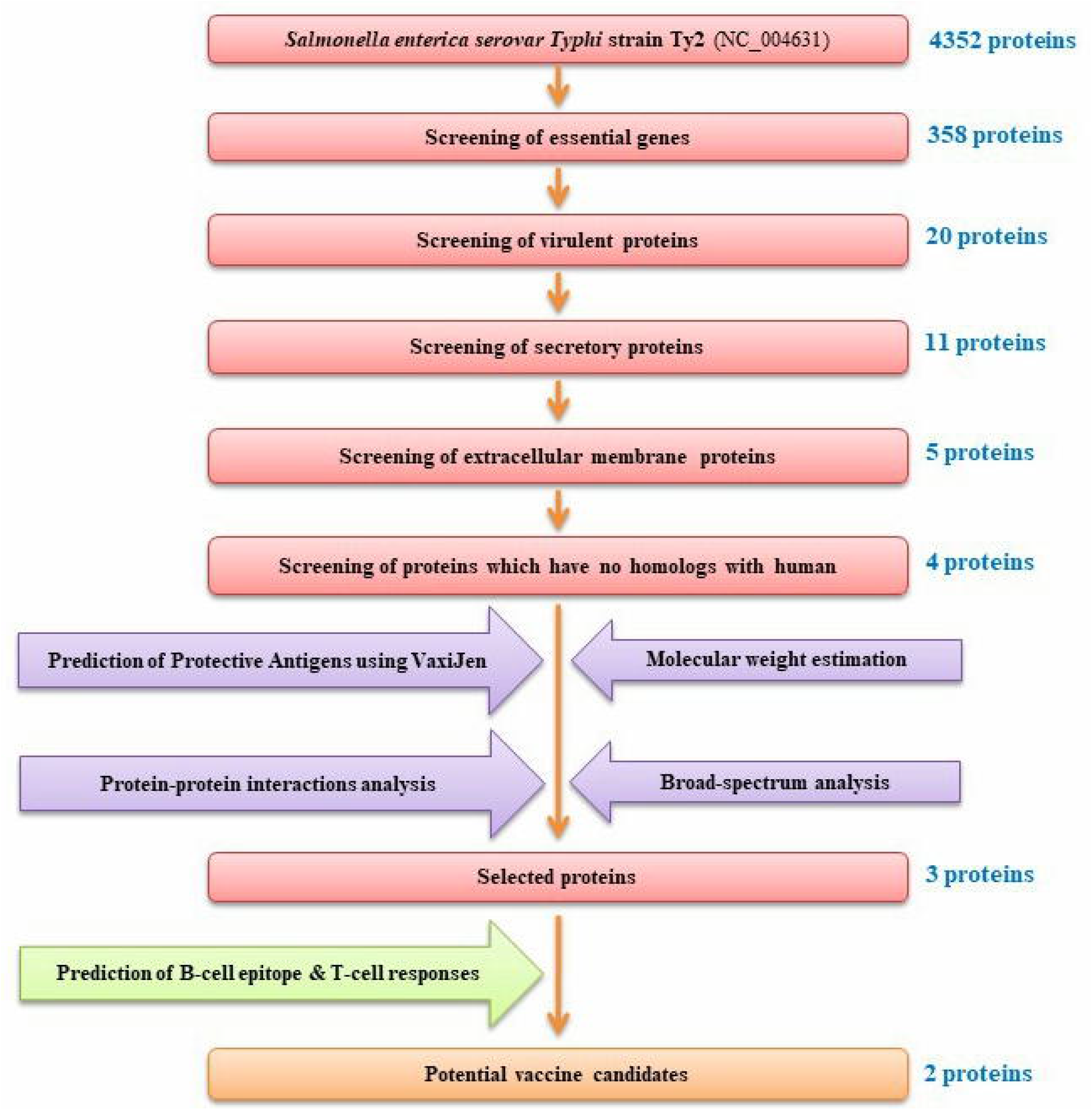
Computational pipelines based on reverse vaccinology and subtractive genomics based approach for screening the potential vaccine targets from *Salmonella typhi*.

**Figure 2:**
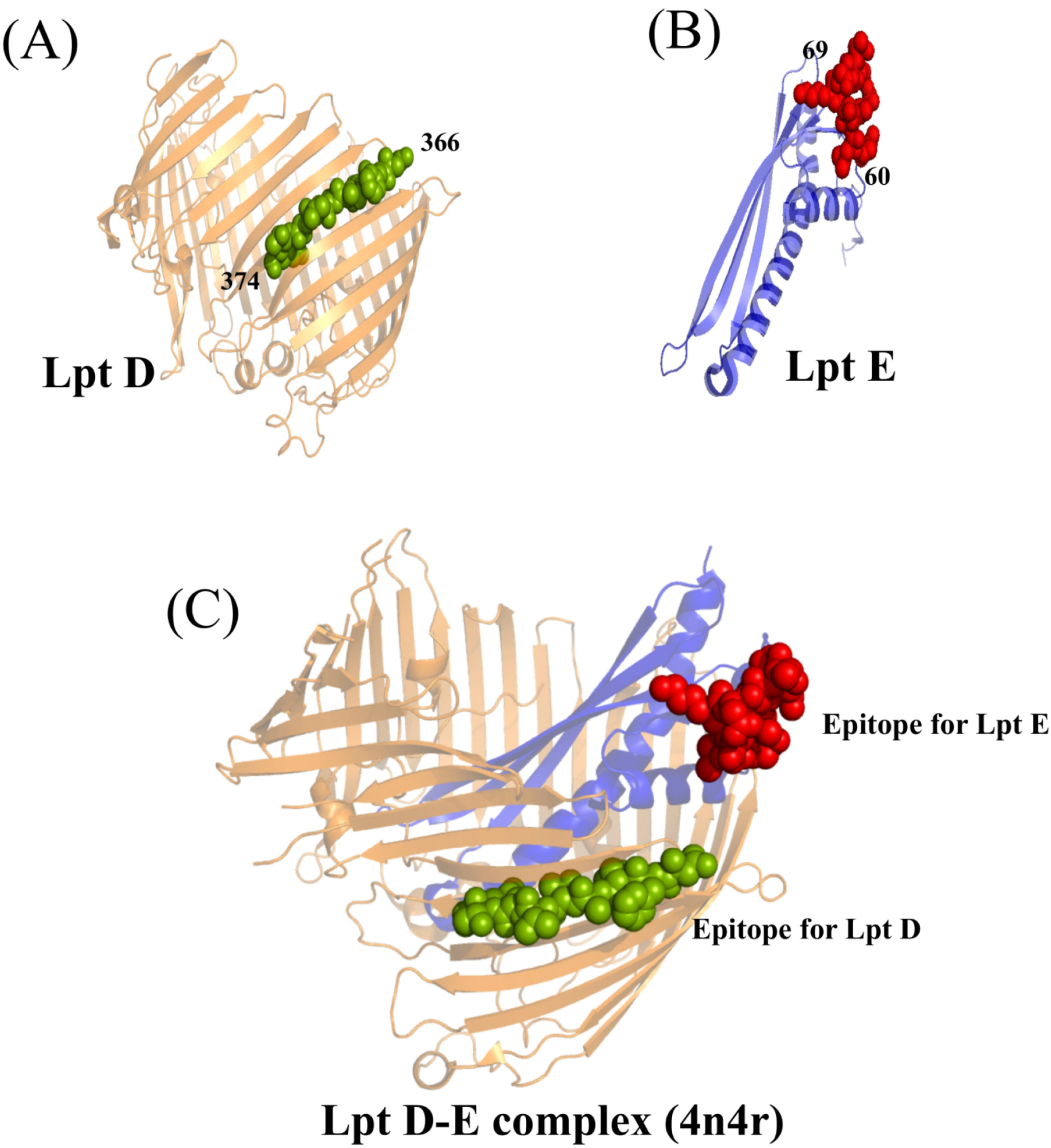
Structural mapping of the epitopes on the PDB structure of LptD-LptE complex. (A) Structure of LptD. The green spheres denote the epitope part of LptD. The residues mapped are from 366 to 374. (B) Structure of LptE. The red spheres denote the epitope part of Lpt E. The residues mapped are from 69 to 74. (C) Structure of LptDE complex (PDB id: 4n4r). The epitopes are mapped for LptD and LptE on the the structure with green and red respectively.

**Table 1:** Details of potential antigenic epitopes that can recognize B cells and T cells

| Name of the Protein | B-cell epitopes predicted by Lbtope | % Probabilty of correct prediction | VaxiJen score | VirulentPred score | CTL epitopes predicted by CTLpred consensus approach | MHC allele (predicted by nHLApred) | Position |
| --- | --- | --- | --- | --- | --- | --- | --- |
| LptD | NFDATVSTKQFQVFN | 81.98 | 0.9712 | 1.0581 | NFDATVSTK | HLA-Cw*0401, H2-Db, H2-Dd, H2-Kb, H2-Kd, H2-Ld, HLA-G, H-2Qa, Mamu-A*01 | 366-374 |
| LptE | NNVNLLDKDTTRKDV | 81.66 | 0.9274 |  | NLLDKDTTR | HLA-Cw*0401, H2-Db, H2-Dd, H2-Kb, H2-Kd, H2-Ld, HLA-G, H-2Qa, Mamu-A*01 | 60-66 |

## 4. Discussion

Lipopolysaccharides (LPS) are unique glycolipids that contribute to permeability barrier properties of the outer membrane of Gram-negative bacteria, thus helping it to survive in the harsh environment and evading attacks from the host immune system [43]. LPS also forms a permeation barrier that inhibits the hydrophobic antibiotics from entering the bacteria which promotes the antibiotic resistance [43][44]. LPS biogenesis is a complex process which are responsible for the transport of LPS across the cellular envelope and the assembly of LPS at the cell surface [45]. Transport of LPS to the outer membrane needs different transporter proteins named Lipopolysaccharide Transporter (Lpt) proteins. The Lpt system contains seven components, the soluble cytoplasmic protein LptB, inner membrane LptF, and LptG, periplasmic LptC anchored to the Inner Membrane by a transmembrane helix, soluble periplasmic LptA, periplasmic LptE anchored to the Outer Membrane by a lipid anchor attached to Cys19 and Outer membrane localized LptD (Figure 3). All of the seven Lpt proteins are essential for the LPS transport to the outer membrane [46]. Two proteins among the group form a plug and barrel conformation in which LptE is located inside the barrel of LptD, which is important for the transport of LPS across the membrane. The primary function of LptDE complex is to place the LPS in the Outer Membrane. According to the current model of LptDE function, LPS arrives from the Inner membrane to the periplasm at the periplasmic N-terminal domain of LptD, causing a conformational change of LptDE [47]. This enables LPS molecules to enter the interior of the barrel. LPS can then move through the lumen of LptD and hence selectively passing through the lateral opening of LptD into the extracellular leaflet of Outer Membrane [47]. Recent studies demonstrated that LptD have the strong ability to generate the immune responses against Vibrio infection and showed a significant reduction of bacterial growth and LPS level, and increased susceptibility to antibiotics which supports that LptD could be the promising target for the development of effective vaccines [48]. Our finding highlighted that LptD, and LptE both are the outer membrane protein responsible for final transport of LPS to the outer leaflet, was widely distributed and highly conserved, and possesses shared surface epitopes among the several major Salmonella pathogens. Both of these extracellular membrane proteins are essential and involved in the secretory pathway, containing the virulence factors and do not have any human homologous, and possess antigenic epitopes. Therefore, both of these proteins have most of the important signatures of being potential vaccines. Thus, LptD and LptE could be the prospective vaccine antigen and therapeutic target for the prevention and control of *Salmonella* infection. Further experimental studies based on our findings would facilitate the development of new vaccines against *S typhi* in near future. Moreover, the reverse vaccinology and subtractive genomics based computational approaches used in this study can also be used for screening the genomes of different pathogens to detect the potential vaccine targets.

**Figure 3:**
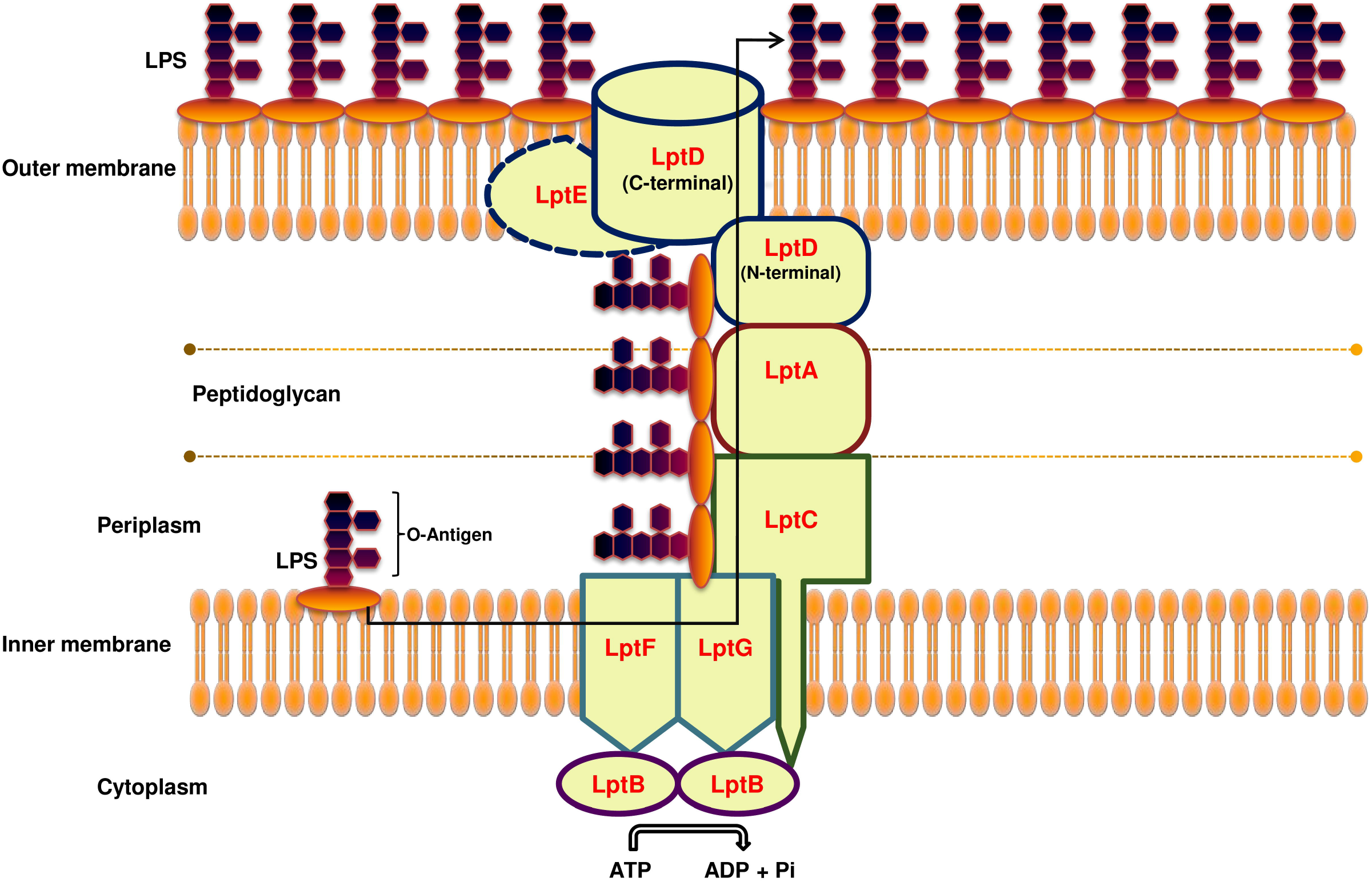
LPS transport system. LptDE complex is essential for the final transport of LPS across the membrane.

## Acknowledgments

SM and KG are thankful to the IISER Kolkata for fellowship support.

## List of abbreviations

LPS: Lipopolysaccharide
*S typhi*: *Salmonella enterica serovar typhi*
BLAST: Basic Local Alignment Search Tool

